# Sequence Analysis for SNP Detection and Phylogenetic Reconstruction of SARS-CoV-2 Isolated from Nigerian COVID-19 Cases

**DOI:** 10.1101/2020.09.25.310078

**Authors:** Idowu A. Taiwo, Nike Adeleye, Fatimah O. Anwoju, Adeyemi Adeyinka, Ijeoma C. Uzoma, Taiwo T. Bankole

## Abstract

**Background:** Coronaviruses are a group of viruses that belong to the Family Coronaviridae, Genus *Betacoronavirus*. In December 2019, a new coronavirus disease (COVID-19) characterized by severe respiratory symptoms was discovered. The causative pathogen was a novel coronavirus known as 2019-nCoV and later as SARS-CoV-2. Within two months of its discovery, COVID-19 became a pandemic causing widespread morbidity and mortality.

**Methodology:** Whole genome sequence data of SARS-CoV-2 isolated from Nigerian COVID-19 cases were retrieved by downloading from GISAID database. A total of 18 sequences that satisfied quality assurance (length ≥ 29700 nts and number of unknown bases denoted as ‘N’ ≤ 5%) were used for the study. Multiple sequence alignment (MSA) was done in MAFFT (Version 7.471) while SNP calling was implemented in DnaSP (Version 6.12.03) respectively and then visualized in Jalview (Version 2.11.1.0). Phylogenetic analysis was with MEGA X software.

**Results:** Nigerian SARS-CoV-2 had 99.9% genomic similarity with four large conserved genomic regions. A total of 66 SNPs were identified out of which 31 were informative. Nucleotide diversity assessment gave Pi = 0.00048 and average SNP frequency of 2.22 SNPs per 1000 nts. Non-coding genomic regions particularly 5’UTR and 3’UTR had a SNP density of 3.77 and 35.4 respectively. The region with the highest SNP density was ORF10 with a frequency of 8.55 SNPs/1000 nts). Majority (72.2%) of viruses in Nigeria are of L lineage with preponderance of D614G mutation which accounted for 11 (61.1%) out of the 18 viral sequences. Nigeria SARS-CoV-2 revealed 3 major clades namely Oyo, Ekiti and Osun on a maximum likelihood phylogenetic tree.

**Conclusion and Recommendation:** Nigerian SARS-CoV-2 reveals high mutation rate together with preponderance of L lineage and D614G mutants. Implication of these mutations for SARS-CoV-2 virulence and the need for more aggressive testing and treatment of COVID-19 in Nigeria is discussed. Additionally, attempt to produce testing kits for COVID-19 in Nigeria should consider the conserved regions identified in this study. Strict adherence to COVID-19 preventive measure is recommended in view of Nigerian SARS-CoV-2 phylogenetic clustering pattern, which suggests intensive community transmission possibly rooted in communal culture characteristic of many ethnicities in Nigeria.

## Introduction

Coronaviruses are a group of viruses that belong to the Family Coronaviridae, Genus *Betacoronavirus* [1, 2]. These viruses are of special interest because they possess the largest genome among RNA viruses and also have the capability to infect a wide host range causing intestinal and respiratory infections in animals and humans [3, 4]. In recent times, coronaviruses have attracted renewed interest in view of a novel coronavirus disease outbreak of 2019 (COVID-19) that originated in Wuhan, China [5]. The causative agent was found to be a novel coronavirus (2019-nCoV) that was identified in December, 2019. Barely two months after its discovery, had the disease become a pandemic of global concern causing widespread morbidity and mortality [6, 7]. Because COVID-19 is a serious respiratory disease, the causative pathogen was later known as severe acute respiratory syndrome coronavirus 2 (SARS-CoV-2).

Before the origin of SARS-CoV-2, six coronaviruses were known to infect man [5, 8]. Out of these, severe acute respiratory syndrome coronavirus (SARS-CoV) and Middle East respiratory syndrome coronavirus (MERS-CoV) caused severe respiratory illness in humans [9, 10]. In 2002, SARS-CoV, originating from Guangdong Province, Southern China, and MERS-CoV emerging from Saudi Arabia caused serious epidemics in 2002/2003 and 2012 respectively [11, 12]. The severity of the respiratory diseases occasioned by these viruses and the resulting wide spread morbidity and mortality had been reported elsewhere [13].

Immediately after its identification, scientists in several laboratories swiftly sequenced SARS-CoV-2 genome making the complete and partial sequences available in public databases. Within few months, the number of viral genome entries in databanks grew exponentially, and this afforded scientists unprecedented opportunity for undertaking molecular, phylogenetic and epidemiological studies on SARS-CoV-2. It was re-established that coronaviruses including SARS-CoV-2 possess the largest genome ranging from 26.4 to 31.7 kb among all RNA viruses [14, 15]. Like other coronaviruses, SARS-CoV-2 contains a positive-sense single-stranded RNA (ssRNA). The first two-third portion of the genome beginning from the 5’ end codes for a polyprotein pp1ab, whose cleavage gave several non-structural proteins (nsps) that are mainly associated with genome replication and transcription [15, 16]. On the remaining portion of coronavirus genome (i.e. towards the 3’ end), there are four genes that encode viral structural proteins namely Spike, (S), Envelope (E), Membrane (M) and Nucleocapsid (N) necessary for viral infectivity and transmissibility [17]. Out of these four structural genes, the S gene has attracted the greatest attention because it encodes the spike protein which binds human angiotensin-converting enzyme (ACE2) for viral attachment and entry into human cells. SARS-CoV-2 appears to be highly optimized for binding to human ACE2 receptor using the receptor binding domain (RBD) on the spike glycoprotein [18].

The first coronavirus genome sequence report in Africa was from a Nigerian index case who travelled to the country from Italy in February, 2020 [19]. In early March, 2020, complete (29,798 nt) and partial (486 nt) genome sequences of SARS-CoV-2 collected from the index case were produced by Nigerian scientists at African Centre of Excellence for the Genomics of Infectious Disease (ACEGID) at the Redeemers University of Nigeria (RUN), and the Centre for Virology and Genomics at the Nigerian Institute of Medical Research (NIMR), Nigeria, respectively. The partial and the complete sequences were deposited in the National Center for Biotechnology Information (NCBI) GenBank (Accession No: MT159778) and the Global Initiative on Sharing All Influenza Data (GISAID) databases (EPI_ISL_413550) respectively. At the time of writing this paper on the 5^th^ August, 2020, additional 41 complete SARS-CoV-2 Nigerian SARS-CoV-2 genome sequences have been added to GISAID by ACEGID to give a total of 42 complete sequence records. Preliminary molecular and phylogenetic analyses gave a phylogenetic tree where the Nigerian index case clustered with a European clade [19]. Subsequent analysis of 18 complete sequences suggested multiple introduction and community transfer in Nigeria [20].

These preliminary reports focused on SARS-CoV-2 spike protein mutations without a comprehensive molecular characterization in terms of number and distribution of SNPs, haplotype, and conserved genomic regions of Nigerian SARS-CoV-2. We hereby present a report of a detailed computational analysis and phylogenetic reconstruction of the Nigerian SARS-CoV-2 using genome data deposited in GISAID database. The results of this study will aid the understanding of variability and evolution of SARS-CoV-2 genome in Nigerian population bearing in mind that such knowledge is crucial for efficient genetic testing, diagnosis, prognosis, and treatment of COVID-19 interventions in Nigeria.

## Materials and Methods

### Data Retrieval

Whole genome sequence data of SARS-CoV-2 isolated from Nigerian COVID-19 cases were retrieved by downloading from GISAID (Global Initiative on Sharing All Influenza Data) database. A total of 43 sequences were deposited as at 18^th^ July, 2020. Out of these, 42 sequences were complete genome sequences when filtered out by selecting the ‘complete’ filter button in GISAID database page implying that one sequence was partial. To ensure acceptable size and quality of SARS-CoV-2 genome sequence for the study, ‘high coverage’ filtering button was also selected. This and quality check by visualization left us with 18 sequences that satisfied all the selection criteria (Table 1). According to GISAID database, ‘complete’ genome sequences are sequences with >29,000 nucleotides (nts) while ‘high coverage’ sequences are those with <1% Ns and no insertions and deletions unless verified by the submitter. The sequences were retrieved by downloading in FASTA file format from GISAID database for further analysis.

**Table 1:** Summary Statistics of the Retrieved Sequences.

| Statistical Parameter | Value |
| --- | --- |
| Mean (nts) | 29863 |
| SD | 35.11 |
| Maximum Size (nts) | 29903 |
| Minimum Size (nts) | 29787 |
| Coeff of Variation (%) | 0.12 |

### Sequence Homology and Mapping

To ensure true homology, non-biological differences (i.e. differences due to technical variations) were removed from retrieved sequences. The sequences were aligned in MAFFT and trimmed at the 5’ and 3’ ends in MEGA X to obtain homologous sequences of 29,787 nts each. Alignment revealed that 49 nucleotides should be removed at the 5’ end while 67 nucleotides should be pruned away at the 3’ end to obtain the 29,787 nts for each of the homologous sequences used for the study. The trimmed sequences were mapped onto a reference genome sequence of SARS-CoV-2 retrieved by downloading from NCBI GenBank (Accession No: NC_045512.2) in order to determine the location of sites on the sampled genome sequences [15]. The 5’ and the 3’ ends of the aligned sequences are shown in Fig. 1.

**Fig 1:**
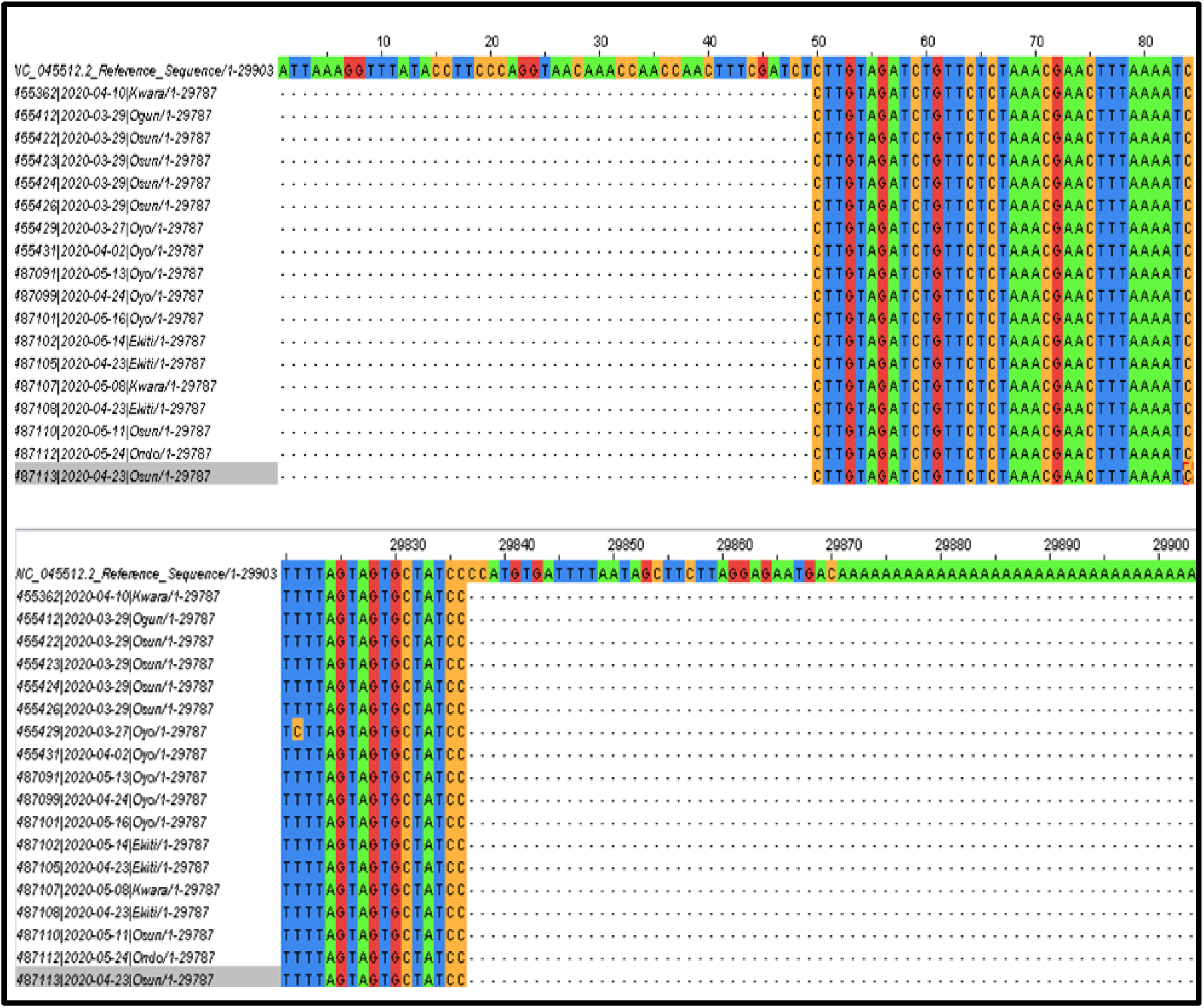
Aligned Sequences Trimmed at 5’ and 3’ ends to Obtain True Homology.

### Multiple Sequence Alignment

Multiple sequence alignment (MSA) was by MAFFT Version 7.471 while phylogenetic reconstruction was by p-distance (in the units of number of base differences per site) was implemented in MEGA X. The genomes were initially aligned with MAUVE to check for large scale genomic changes including large deletions, gene inversion, and genome rearrangements. Then, the sequences were re-aligned in MAFFT (Fig.1) to produce aligned sequences that were fed into DnaSP for SNP and haplotype analysis and subsequently into Jalview 2.11.1.0 for visualization and automatic determination of allelic frequency of SNPs.

### Variation and SNP Analysis

Genomic locations of SNPs were determined relative to the SARS-CoV-2 reference genome sequence obtained from NCBI GenBank (Accession No: NC 045512.2). A polymorphic site is considered as a genomic location with the frequency of the second most frequent allele greater than 1%; otherwise, they were referred to as monomorphic. Rare alleles are those with minor allele frequency (MAF) less than 1%, and the viruses carrying them were rare variants. Detection of SNPs, linkage disequilibrium and haplotype analysis was done in DnaSPv6.12.03 while SNP distribution and density was obtained by a series of MS Excel formulae packaged by us.

### Phylogenetic Analysis

Maximum likelihood phylogenetic tree construction was implemented in MEGA X using sequences that had been aligned by MAFFT employing Tamura-Nei evolutionary model under assumption of uniform nucleotide substitution [21, 22]. The tree that had topology with superior log likelihood value was selected from the initial trees by applying neighbor-joining (NJ) and bio-NJ algorithm in a heuristic search. Analysis of the tree was based on topology and clustering pattern analysis. Tree validation was by 100 bootstrap replicates [23].

## Results

### Summary Statistics of the Retrieved Sequences

The summary statistics of 18 SARS-CoV-2 complete sequences that satisfied our selection criteria are presented in Table 1. The complete genome sequences of mean size 29863 ± 35.11 (mean+SD) were obtained from the retrieved sequences from SARS-CoV-2 isolated from 18 subjects (8M: 6F: 4 Unknown Sex) from 6 states in Nigeria namely, Kwara, Ogun, Osun, Oyo, Ekiti and Ondo.

### Results of SNP Analysis

Nigerian SARS-CoV-2 had 99.9% genomic similarity with four large conserved genomic regions (Region 1: 13020-14407, Region 2: 16742-18876, Region 3: 20375-21897, and Region 4: 23407-25470). A total of 66 SNPs were identified in the SARS-CoV-2 genomes used for this study out of which 31 were informative. All the SNPs were diallelic except a SNP located at 29791 nt which was triallelic. The overall nucleotide diversity among the SARS-CoV-2 genomes analyzed was Pi = 0.00048. The average frequency of SNPs on SARS-CoV-2 genomes in Nigeria was 2.22 SNPs per 1000 nts (Table 2). The SNPs were not, however, evenly distributed in the entire genome: highest SNP densities were observed in the 5’UTR and the 3’UTR regions with SNP frequency of 3.77 and 35.4 respectively. The coding regions with the highest SNP density was ORF10 (8.55 SNPs/1000 nts). No SNP was detected in ORFs 6, 7a, 7b and E gene.

**Table 2:**
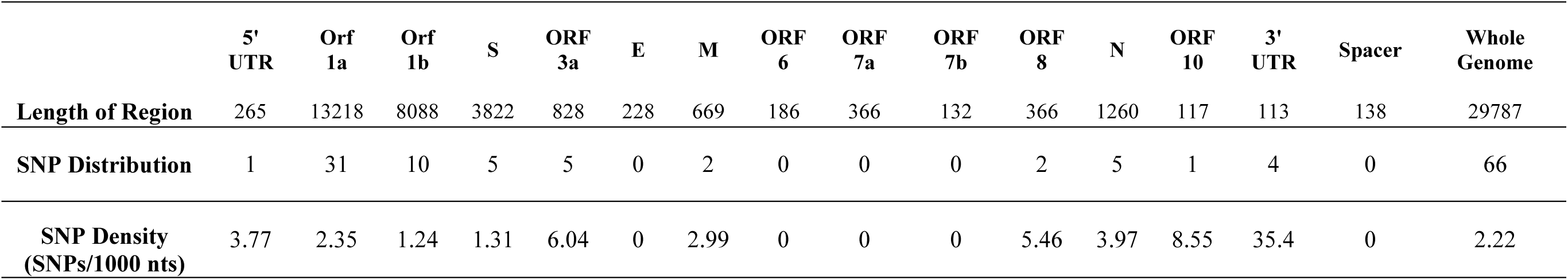
Distribution and Density of SNPs in SARS-CoV-2 Isolated in Nigeria.

|  | **5' UTR** | **Orf 1a** | **Orf 1b** | **S** | **ORF 3a** | **E** | **M** | **ORF 6** | **ORF 7a** | **ORF 7b** | **ORF 8** | **N** | **ORF 10** | **3' UTR** | **Spacer** | **Whole Genome** |
| --- | --- | --- | --- | --- | --- | --- | --- | --- | --- | --- | --- | --- | --- | --- | --- | --- |
| **Length of Region** | 265 | 13218 | 8088 | 3822 | 828 | 228 | 669 | 186 | 366 | 132 | 366 | 1260 | 117 | 113 | 138 | 29787 |
| **SNP Distribution** | 1 | 31 | 10 | 5 | 5 | 0 | 2 | 0 | 0 | 0 | 2 | 5 | 1 | 4 | 0 | 66 |
| **SNP Density**<br>**(SNPs/1000 nts)** | 3.77 | 2.35 | 1.24 | 1.31 | 6.04 | 0 | 2.99 | 0 | 0 | 0 | 5.46 | 3.97 | 8.55 | 35.4 | 0 | 2.22 |

### Haplotype Analysis for Delineation of L and S Lineages

Linkage analysis involving pairwise comparison of SNPs showed that several SNPs are in 2-locus linkage disequilibrium (LD) in SARS-CoV-2 genome with majority of the SNPs above the significant (P<0.05) horizontal threshold line indicated in Fig. 2. The total number of significant pairwise haplotypes detected by Fisher’s exact test was 108 out of which 12 were highly significant (P<0.001) by Bonferoni procedure (Table 3). Two linked SNPs: a C/T SNP at location 8782 nt in the orf1ab region and a T/C SNP at 28144 nt in ORF8 region identified by Cui *et al*., [1] were absolutely linked in this study (Table 4). The frequency of CT haplotype (L lineage) and TC (S lineage) were 13 (72.2%) and 5 (27.8%) respectively (Fig. 3).

**Table 3:** Haplotypes Obtained from Sites in Highly Significant Linkage.

|  | Site 1 | Site 2 | Distance (nts) | Linkage Diseq. (D) |
| --- | --- | --- | --- | --- |
| 1 | 241 | 3037 | 2796 | 0.238*** |
| 2 | 241 | 14408 | 14167 | 0.238*** |
| 3 | 241 | 23403 | 23162 | 0.238*** |
| 4 | 3037 | 14408 | 11371 | 0.238*** |
| 5 | 3037 | 23403 | 20366 | 0.238*** |
| 6 | 8782 | 22468 | 13686 | 0.201*** |
| 7 | 8782 | 28144 | 19362 | 0.201*** |
| 8 | 8782 | 28878 | 20096 | 0.201*** |
| 9 | 14408 | 23403 | 8995 | 0.238*** |
| 10 | 22468 | 28144 | 5676 | 0.201*** |
| 11 | 22468 | 28878 | 6410 | 0.201*** |
| 12 | 28144 | 28878 | 734 | 0.201*** |

\*\*\*P < 0.001 (Bonferoni Correction)

**Table 4:** Haplotype Analysis for the Frequency of L and S Lineages.

| Site 8782 | Site 28144 | Haplotype | Lineages | Number | Frequency (%) |
| --- | --- | --- | --- | --- | --- |
| C | T | CT | L | 13 | 72.2 |
| T | C | TC | S | 5 | 29.8 |

**Fig 2:**
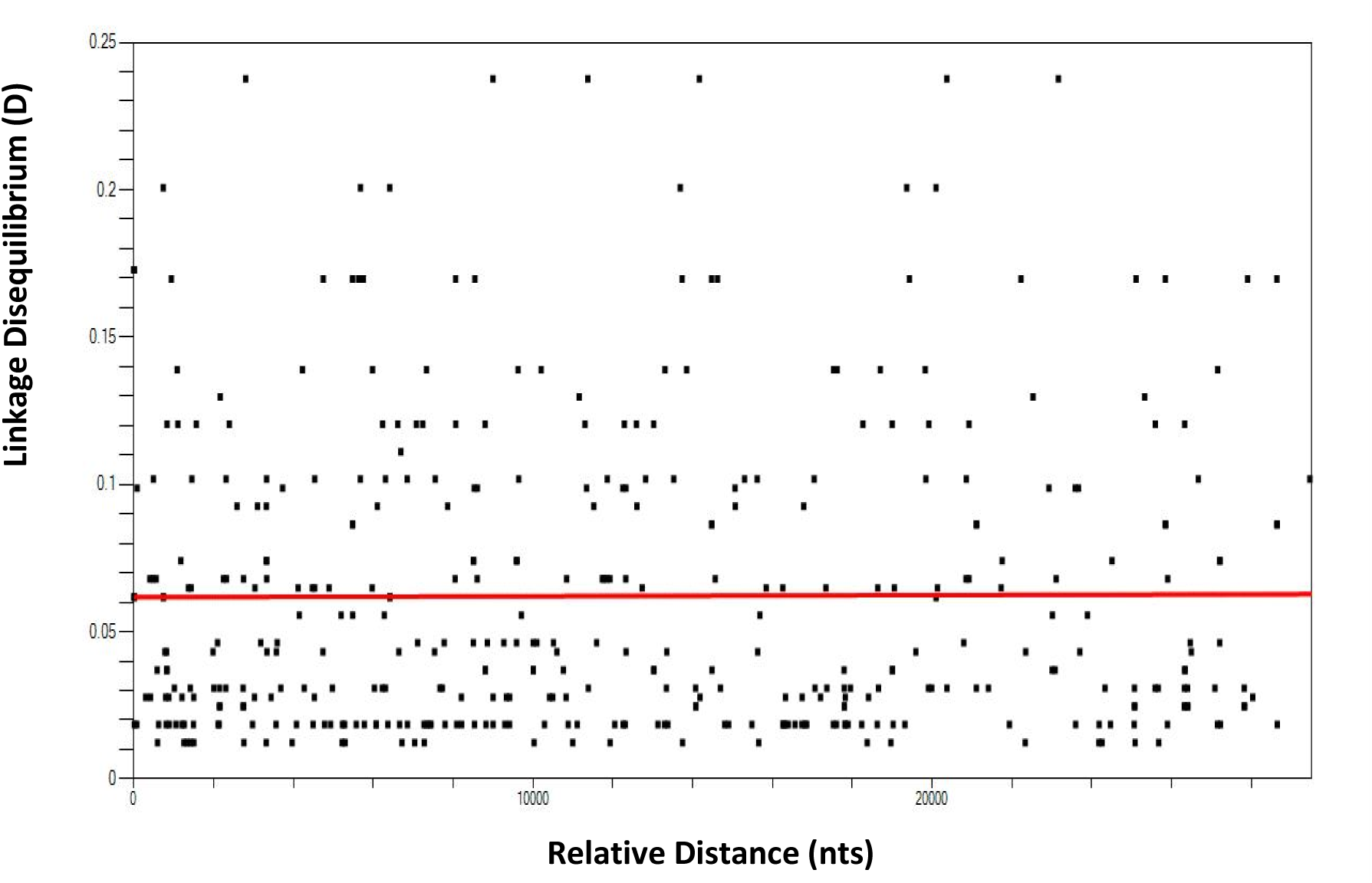
SNPs in Linkage Disequilibrium in Nigerian SARS-CoV-2 Genome. **Note:** SNPs above the line are in significant linkage disequilibrium

**Fig 3:**
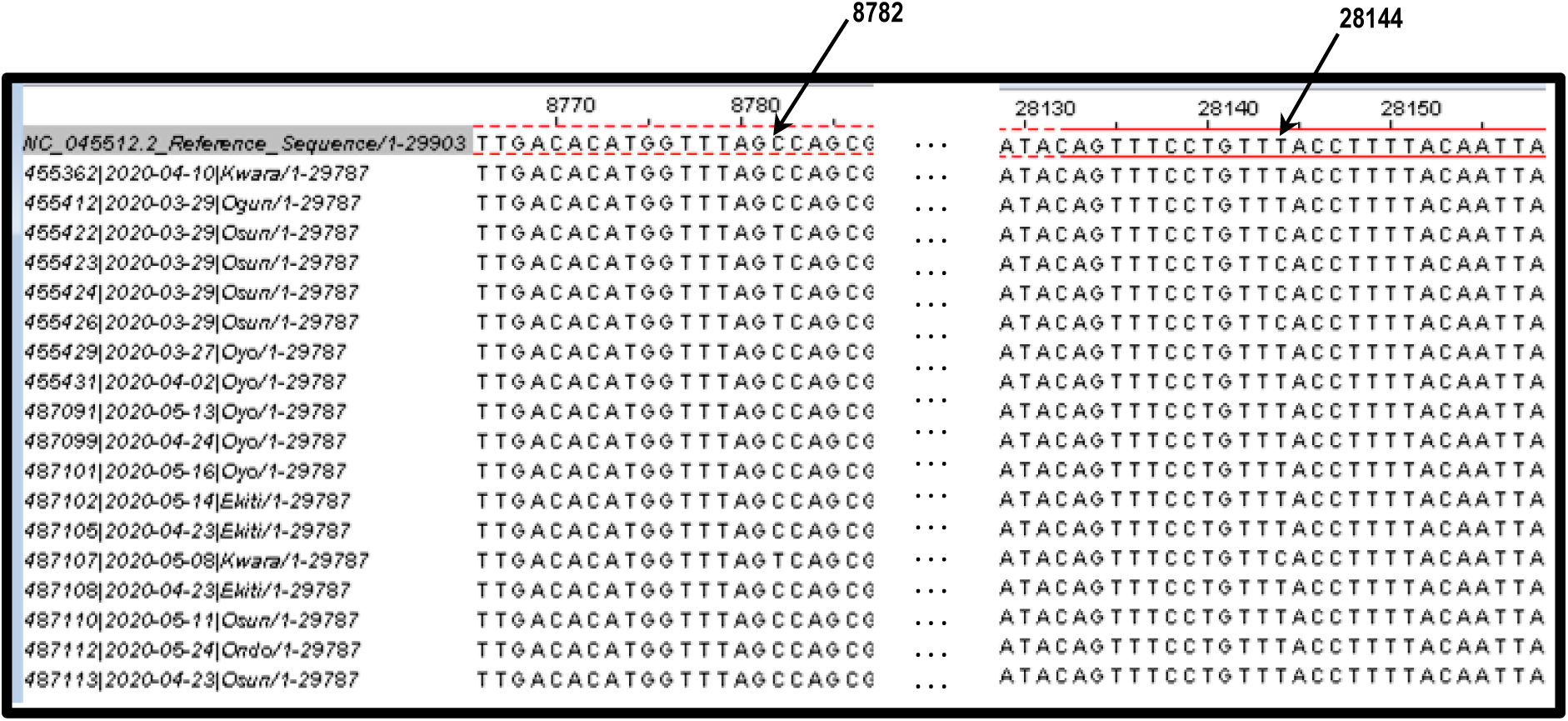
Complete Linkage Disequilibrium of C8782 and T28144 Locations in the Genome of Nigerian SARS-CoV-2 Genome.

### Results of D614G Mutation Analysis

Multiple sequence alignment of translated SARS-CoV-2 spike (S) gene revealed a D/G amino acid substitution at position 614 of the spike protein (Fig. 4). There was a preponderance of D614G mutation because majority 11 (61.1%) of the analyzed genomic sequences of SARS-CoV-2 had G at 614 position of spike protein as compared to 7 (39.9%) that had D at the site.

**Fig 4:**
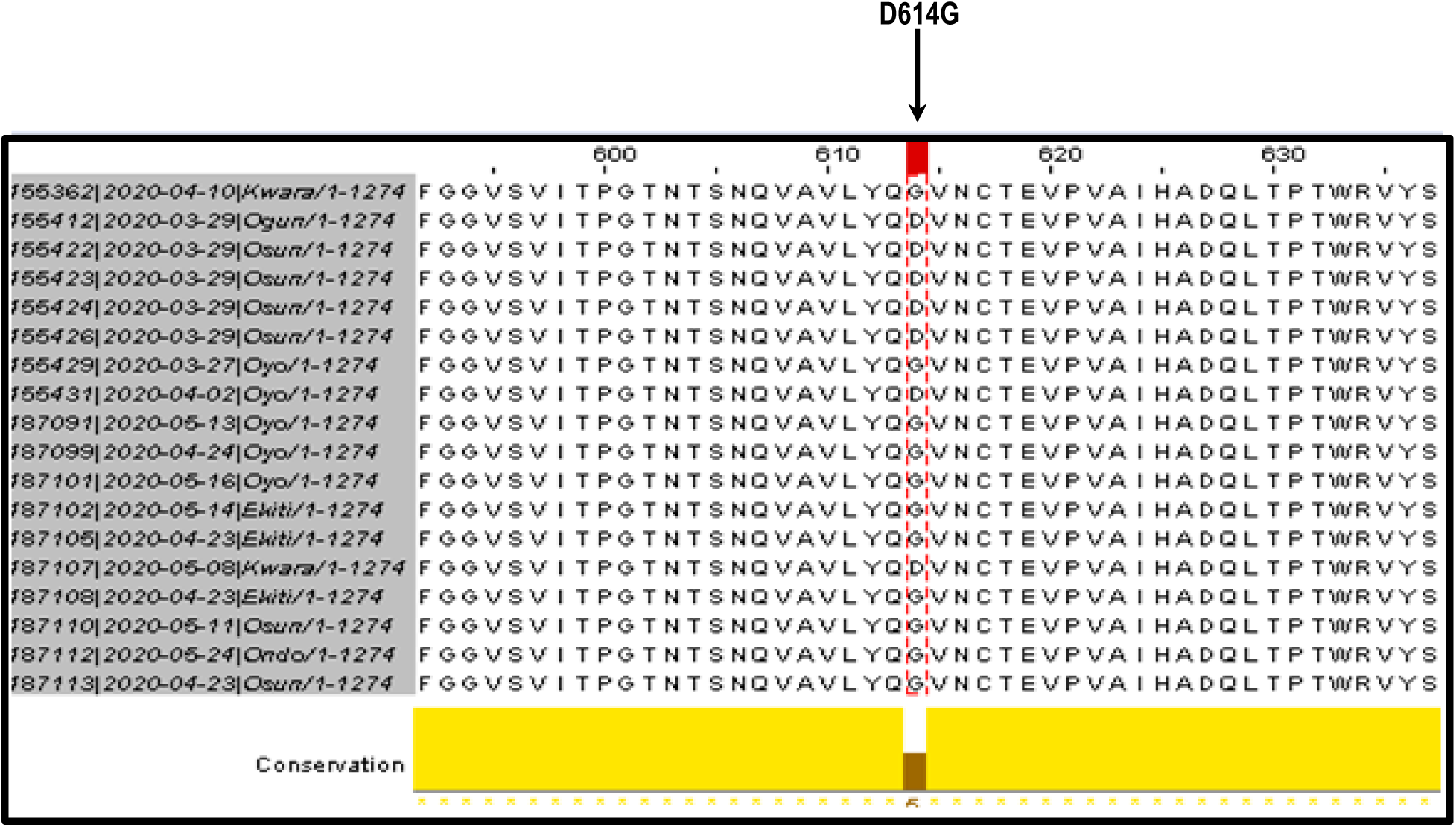
Multiple Sequence Alignment of Translated Spike Protein Revealing Preponderance of D614G Mutation.

**Fig 5:**
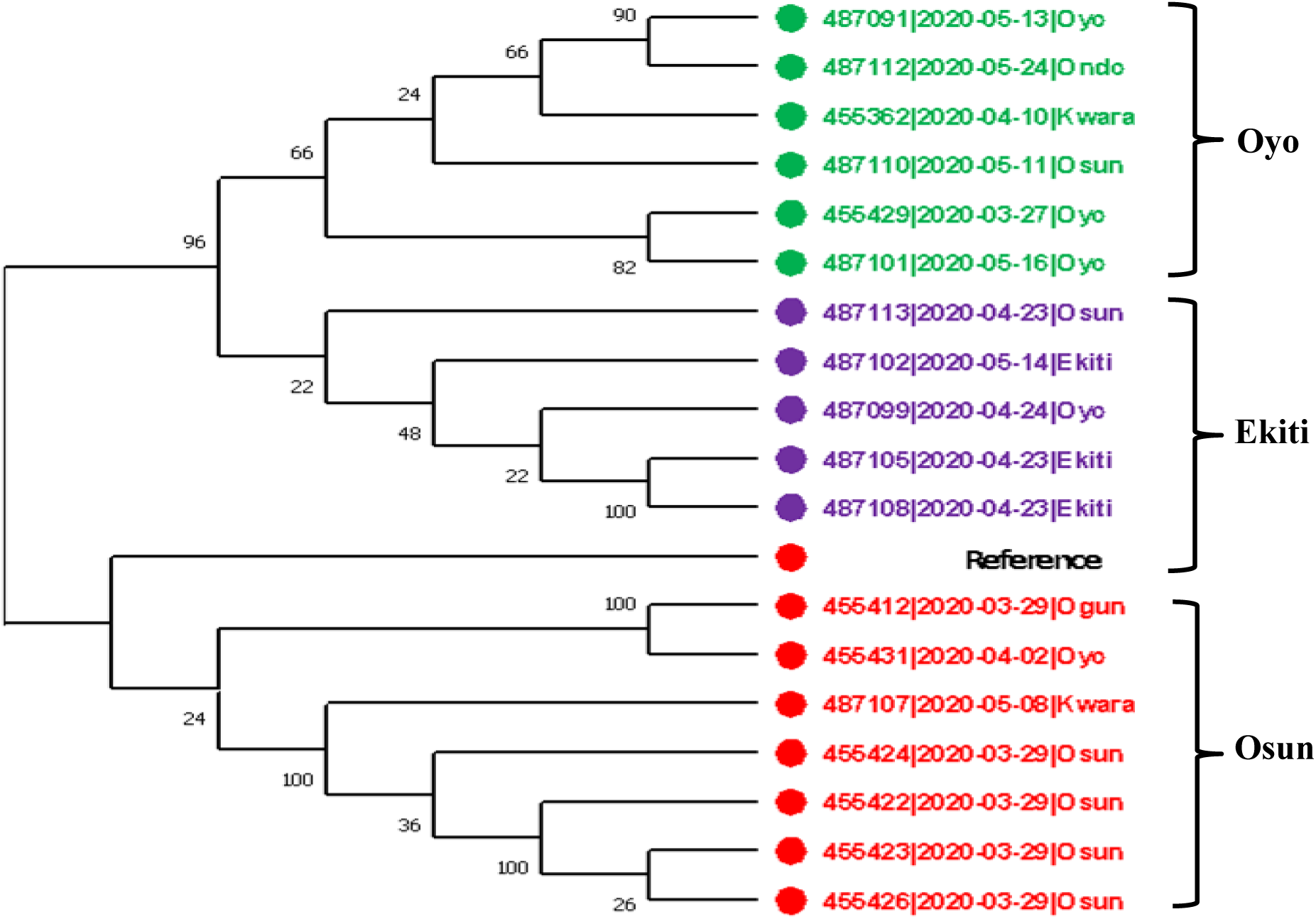
Maximum Likelihood Phylogenetic Tree of Nigerian SARS-CoV-2 Using Whole Genome Sequences.

### Phylogenetic Analysis

Clustering analysis of the maximum likelihood phylogenetic tree gave 3 major clades. On the basis of the preponderance of taxa, the clusters were identified as Oyo, Ekiti and Osun. Reference sequence clustered with Osun clade. Majority of the nodes have high bootstrap support (60%-100%). The node that defines Ekiti clade had the lowest bootstrap support of 22%.

## Discussion

In the present study, whole-genome sequence characterization and phylogenetic reconstruction of SARS-CoV-2 isolated from Nigerian COVID-19 subjects was carried out. Out of the 42 Nigerian complete genome sequences in GISAID database, only 18 fulfilled our selection criteria for inclusion in the study. In view of the implications of quality of results for validity of results, it was better, in our opinion, to use 18 sequences of acceptable quality rather than using 42 sequences where more than half of the sequences are of low or questionable quality.

Nigerian SARS-CoV-2 had 99.9% implying 0.01% dissimilarity in accordance with the global trend [15, 16]. It was therefore not surprising that four large conserved genomic regions were identified in this study. This observation supports the well-established view that the novel virus has a recent origin estimated to be between 6^th^ October and 11^th^ December 2019 [24]. Furthermore, Nigeria has extended family and communal structure, which could favour community transmission of the novel virus. Thus, Nigeria’s socio-cultural attributes may contribute (at least in part) to the high degree of similarity and the ethnic based clustering pattern on the phylogenetic tree obtained in this study. It will be of interest to carry out comparative analysis of conserved regions involving genomes of SARS-CoV-2 in other parts of the world to compare their tree topologies with that of Nigeria.

Despite the high degree of similarity, some degree of diversity resulting from SNPs found in SARS-CoV-2 genome, was detected. The number of SNPs detected in this study was low when compared with other studies [25]. A limitation of this study is the small sample size of eighteen genome sequences used in this study, which we consider too small for detection of all SARS-CoV-2 SNPs in Nigeria. In subsequent studies, it will be of interest to see if distribution of SNPs and conserved regions identified in this study are peculiar to Nigerian SARS-CoV-2 or have complete overlap with genomes of SARS-CoV-2 found elsewhere. If there are regions of non-overlap, any attempt to produce testing kits with high sensitivity for Nigeria’s COVID-19 cases should take note of the conserved regions and SNP distribution in the genomes of SARS-CoV-2 detected in this study.

The frequency of SNPs observed in the Nigerian SARS-CoV-2 genome is higher than twice the frequency of SNPs in human genome which is generally taken to be 1 SNP per 1000 bps. The UTRs, especially 3’UTR, are mutation hot spots in Nigerian SARS-CoV-2 genome. This is expected because UTRs, unlike coding sequences, are generally under more relaxed or neutral selection pressure which allow mutations to accumulate at a higher rate in that region [25]. Relatively high SNP densities were also recorded in some of the coding regions of Nigerian SARS-CoV-2. An example is ORF 10 region with a SNP density of 8.55 SNPs/1000 nts. Although coronaviridae have proofreading capability due to their exonuclease activity during nucleotide replication, mutation rate of SARS-CoV-2 was estimated at ∼6 × 10^− 4^ nucleotides/genome/year with the capacity to mutate during human-to-human transmission [24]. We share the view that characterizing organisms and viruses by haplotypes is more informative than using individual SNPs. Such haplotype analysis gave results which agreed with an early report by Cui *et al*., [1] who proposed two major lineages namely the L lineage carrying CT haplotype at the closely linked positions C8782 and T28144 respectively and the S lineage with nucleotides T and C (TC haplotype) at the corresponding genome locations. The higher frequency of L lineage viruses which was higher than those of S lineage in this study is consistent with the original finding by Cui *et al*., [1]. Few recombinants were observed in their study unlike the present study where both loci were absolutely linked such that no recombinants were recovered. According to Cui *et al*., [1], identification as L and S lineages was based on the amino acids resulting from T28144C substitution in which leucine (L lineage) is replaced with serine (S lineage).

Similar increased frequency of viruses in the L lineage as compared to those in the S lineage was observed at 23403 nt position where A23403G nucleotide substitution caused D614G mutation on SaRS-CoV-2 spike protein. Majority of SARS-CoV-2 in Nigeria were G614 (G type) mutants as opposed to the D614 (D type) ancestral form in agreement with the global trend [26]. Preponderance of L and G variants in many populations was accounted for by their increased Darwinian fitness whereby these newly evolved strains have competitive advantage over their alternative and more ancestral forms in terms of infectivity, transmissibility and, possibly, virulence. Others felt that the preponderance of the L and the G mutants might result from sampling bias and founder effect [27].

Results of studies just concluded in the Center for Genomics Noncommunicable and Personalized Healthcare (CGNPH) in the University of Lagos, Nigeria (not yet published), suggested that SNPs and mutation pattern of SARS-CoV-2 vary between continents and, in some cases, between different countries in the same continent. In view of the fact that testing kits are generally nucleic acid based and the potential of SARS-CoV-2 to evade detection due to its mutational dynamics, it is difficult or impossible to have universal testing kits with equal efficiency for detecting the novel coronavirus in all parts of the world [28]. Therefore, the identified conserved regions and the distribution of SNPs in the genome of SARS-CoV-2 circulating in Nigeria has implications for Nigeria-based intervention programmes in the area of viral detection, drug designing and treatment options as had been generally noted by Vankadari [29]. According to Puty *et al*., [28], it is important for every population to identify key SARS-CoV-2 mutations and SNPs for appropriate population-based intervention specific for its population. The view of Wang *et al*., [15] that limited sensitivity of SARS-CoV-2 detection kits is possibly associated with genetic variation of the virus is pertinent. Thus, any meaningful intervention activity should consider SARS-CoV-2 strains in the concerned population.

## Acknowledgement

We thank scientists at African Centre of Excellence for Genomics of Infectious Disease (ACEGID) at the Redeemers University of Nigeria (RUN), and Centre for Human Virology and Genomics, Nigerian Institute of Medical Research (NIMR), Nigeria, respectively for depositing complete and partial SARS-CoV-2 sequences at GISAID and NCBI GenBank respectively. We also thank their partners and collaborators namely Nigerian Centre for Disease Control (NCDC), Centre for Human and Zoonotic Virology (CHAZVY), College of Medicine, University of Lagos, Nigeria, and Infectious Disease Hospital, Yaba, Nigeria.

